# Scalable preprocessing for sparse scRNA-seq data exploiting prior knowledge

**DOI:** 10.1101/142398

**Authors:** Sumit Mukherjee, Yue Zhang, Joshua Fan, Georg Seelig, Sreeram Kannan

## Abstract

**Motivation:** Single cell RNA-seq (scRNA-seq) data contains a wealth of information which has to be inferred computationally from the observed sequencing reads. As the ability to sequence more cells improves rapidly, existing computational tools suffer from three problems. (1) The decreased reads-per-cell implies a highly sparse sample of the true cellular transcriptome. (2) Many tools simply cannot handle the size of the resulting datasets. (3) Prior biological knowledge such as bulk RNA-seq information of certain cell types or qualitative marker information is not taken into account. Here we present UNCURL, a preprocessing framework based on non-negative matrix factorization for scRNA-seq data, that is able to handle varying sampling distributions, scales to very large cell numbers and can incorporate prior knowledge.

**Results:** We find that preprocessing using UNCURL consistently improves performance of commonly used scRNA-seq tools for clustering, visualization, and lineage estimation, both in the absence and presence of prior knowledge. Finally we demonstrate that UNCURL is extremely scalable and parallelizable, and runs faster than other methods on a scRNA-seq dataset containing 1.3 million cells.

**Availability:** Source code is available at https://github.com/yjzhang/uncurl_python

## 1 INTRODUCTION

High-throughput scRNA-seq technologies (Klein *et al.*, 2015; Rosenberg *et al.*, 2017; Zheng *et al.*, 2017; Grun and van Oudenaarden, 2015) can provide biological insights such as revealing cell type composition (Zeisel *et al.*, 2015; Baron *et al.*, 2016), cell lineage relationships (Trapnell *et al.*, 2014; Shin *et al.*, 2015; Welch *et al.*, 2016; Setty *et al.*, 2016), or even spatial relationships (Satija *et al.*, 2015; Halpern *et al.*, 2017) between cells in heterogeneous multi-cellular systems. Enabling such insights are two key advantages of single cell transcriptomic datasets. First, having information about individual cells helps avoid aggregation and conflation of traits from disjoint groups of cells within a mixed sample (Blyth, 1972). Second, scRNA-seq can generate a very high-dimensional dataset, both in terms of the number of cells and genes that can be assayed, compared to other methods with single-cell resolutions. However, advanced computational methods are required to extract latent biological information from the raw read-counts, which provide only a heavily sampled version of the full cellular transcriptome (Trapnell, 2015; Wagner *et al.*, 2016).

Most commonly used computational tools for cell type identification (Jain and Dubes, 1988; Ding and He, 2004), lineage estimation (Trapnell *et al.*, 2014; Welch *et al.*, 2016; Setty *et al.*, 2016) and similar applications rely on an initial dimensionality reduction step using methods such as PCA (Abdi and Williams, 2010), LLE (Roweis and Saul, 2000) or tSNE (Maaten and Hinton, 2008). However, these algorithms assume that the underlying data is drawn from a Gaussian or a t-distribution, an assumption that does not always hold for scRNA-seq data (Grun *et al.*, 2014). The discrepancy between the assumed and actual distribution fundamentally limits the accuracy of the resulting predictions. In addition to such general purpose preprocessing methods, several tools were developed to specifically deal with scRNA-seq data (Dijk *et al.*, 2017; Wang *et al.*, 2017; Pierson and Yau, 2015). However, these approaches do not scale well with increasing cell number. Finally, all existing methods rely almost exclusively on unsupervised learning and do not incorporate useful and commonly available prior information such as bulk gene expression data or cell type specific marker genes to guide the analysis process.

Here, we introduce UNCURL, a preprocessing framework for scRNA-seq data that addresses these shortcomings by estimating the true transcriptomic state of the cells prior to the sampling effect of RNA-seq. The preprocessed data from UNCURL can then be directly used as input by most major unsupervised learning algorithms commonly used in the context of scRNA-seq data. An overview of the algorithmic workflow of UNCURL can be seen in Fig 1A. The main technical contribution of UNCURL is a generalized non-negative matrix factorization (NMF) that explicitly accounts for the most likely sampling distribution of the dataset. Furthermore, UNCURL incorporates an accelerated optimization method tailored for sparse input data, so as to handle datasets with millions of cells efficiently.

**Fig. 1.**
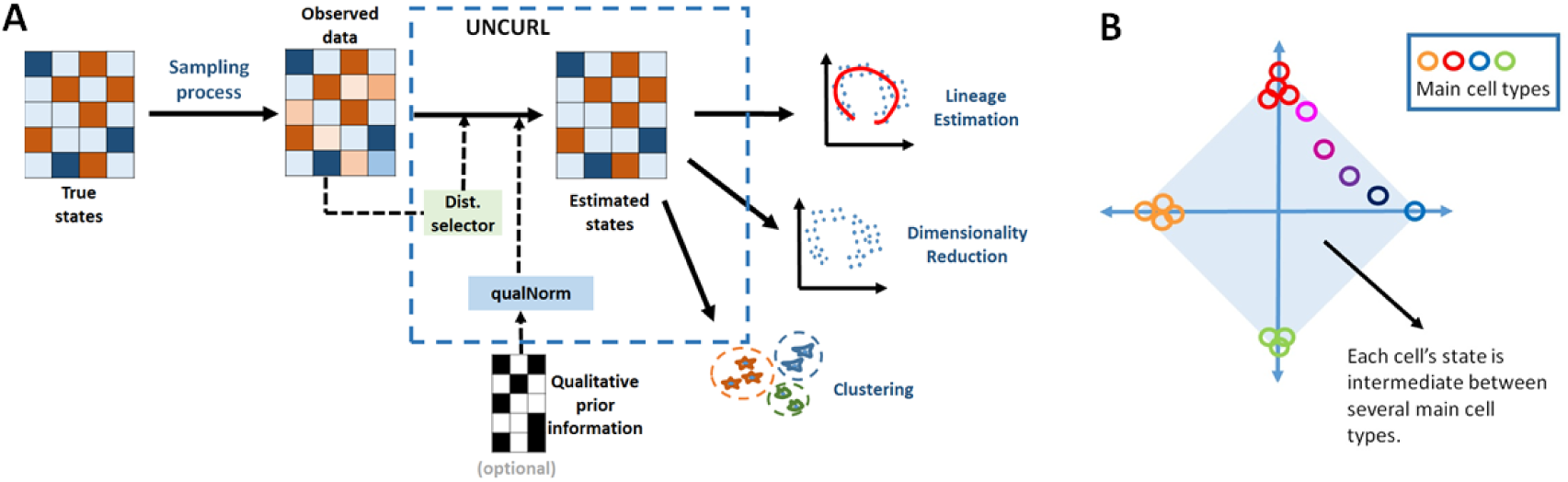
A) The primary input for UNCURL is the highly sampled single cell sequenced data and optionally any prior information that is known about the specific dataset. UNCURL then converts the observed sampled data to an estimated version of the true data using a novel sampling model aware matrix factorization. This can then be used in downstream unsupervised learning tasks. B) The convex mixture of cell states assume that all cell states lie in the convex hull spanned by a few extreme cell types.

Our algorithm exploits the low-dimensional nature of the true biological state matrix, i.e. it assumes that each cell is in a convex combination of a few archetypal cell-states. Under this assumption, the true state matrix can be expressed as a product of an archetypal main state matrix, M, comprising of gene-expression in the archetypal states, and a matrix of mixing coefficients, W, a cluster-by-cell matrix for which each column sums to 1. We demonstrate that working with the estimated (and factorized) true state matrix considerably improves performance of state-of-the-art methods as compared to directly operating on the sequencing data.

Additionally, UNCURL allows for the integration of prior information which leads to large improvements in accuracy. To enable semi-supervised learning, UNCURL’s toolbox contains a method (qualitative normalization, or qualNorm) for standardizing any prior biological information including bulk RNA-seq data, microarray data or even information about individual marker gene expression to a form compatible with scRNA-seq data. We demonstrate that initialization using prior knowledge in an appropriately standardized manner dramatically improves performance compared to unsupervised learning.

## 2 METHODS

### 2.1 State Estimation

#### 2.1.1 Procedure

An implicit assumption shared by many scRNA-seq data analysis tools is that any biological sample contains a limited number of cell types and that any individual cell can be considered a mixture of these cells. Here, we make this convex mixture model explicit, which leads to a model similar to NMF. NMF is classically used when the entries have Gaussian noise (Lee and Seung, 2001) and has been found beneficial in analyzing gene expression data gleaned from microarrays (Brunet *et al.*, 2004). In scRNA-seq, the sequencing process can produce noise following several different distributions such as Gaussian, log-normal, Poisson and Negative Binomial, potentially with zero-inflation (Pierson and Yau, 2015). While NMF can be directly applied to the scRNA-seq data (Shao and Hofer, 2017), utilizing the sampling distribution becomes critical especially when the number of reads per cell is small. While the sampling distribution is carefully modeled in differential expression studies (Anders and Huber, 2010), the most commonly used algorithms for visualization, cell-type identification as well as lineage estimation do not account for this model. Thus, while factoring the matrix, we need to account for the sampling distribution in order to estimate the true cell-state matrix and mixing coefficients accurately from the observed gene expression matrix.

We assume that we are provided with a data matrix *X ∈* ℝ^*n×d*^, where *n* is the number of genes and *d* is the number of cells, and *k*, the number of cell types. Let *X*_*g,c*_ denote the count measured for gene *g* in cell *c*, and let 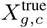 be the relative abundance of gene *g* in cell *c*. We assume that the true matrix has a non-negative decomposition into two factors, i.e., *X*^true^ = *M × W*. Here, *M ∈* ℝ^*n×k*^ is the matrix of cell type means of the *k*-archetypal states with *M*_*g,j*_ denoting the expression of gene *g* in archetype *k*. *W ∈* ℝ^*k,d*^ is the mixture parameter matrix which stores each cell as a convex combination of the archetypes, i.e., *W*_*j,c*_ is the contribution of archetype *j* to cell *c*. Thus *W* satisfies **1**^*k*^*W*_*j,c*_ = 1 and *W*_*j,c*_ *≥* 0.

Since we do not observe directly the relative abundance but only a sampled version, we assume that there is a channel that connects the true abundance to the observed abundance, ℙ(*x | x*^true^, *θ*), i.e, given a value of *x*^true^ there is a certain distribution on *x* with some parameters *θ*. Note that we have not used a subscript for *g* and *c* to emphasize that the same distribution is used for all genes. We give three examples here, but our framework works with general distributions: (1) ℙ(*x |x*^true^) is a Gaussian distribution with mean *x*^true^ and a fixed variance 1 (say). (2) ℙ(*x |x*^true^) is a Poisson distribution with mean *x*^true^. (3) ℙ(*x |x*^true^) is such that log(*x*) is a Gaussian with mean log(*x*^true^) and variance 1. Our goal now is to maximize the log-likelihood of the observed data matrix *X*, by finding the optimal *M* and *W*. Note that *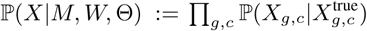.*

This problem is non-convex but the sub-problems of estimating either *M* or *W* with the other matrix fixed are convex problems for many common sampling distributions, including the ones mentioned above. We thus utilize an alternating maximization algorithm to estimate these model parameters as follows:

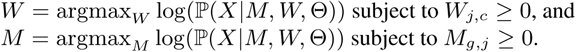

We repeat these two steps iteratively till convergence or till a maximum number of iterations. Once converged, we normalize the columns of *W* to sum to 1 to ensure the condition **1**^*k*^*W*_*j,c*_ = 1 is satisfied. We note that each of the steps is convex for many distributions and can be solved by gradient descent. Unlike the Gaussian and log-normal, Poisson Log-Likelihood does not have closed form solutions for even the sub-problems i.e. identifying *M* and *W*. Hence, Poisson state estimation requires the use of gradient descent based strategies, most of which have sub-linear convergence guarantees only for functions with Lipschitz continuous gradients. This is not true for the Poisson Log-Likelihood. However, (Bauschke *et al.*, 2016) utilized a different definition of smoothness and derived a generalized algorithm (NoLips) that is capable of achieving a sublinear rate of convergence for a class of non-Lipschitz continuous functions including the Poisson Log-Likelihood. Here we use a custom alternating minimization approach using the NoLips algorithm to optimize the Poisson Log-Likelihood with additional modifications to allow for faster computation for sparse matrices and ability to parallelize the computation (see supplementary materials).

#### 2.1.2 State estimation for different distributions

While UNCURL can easily be extended to different sampling distributions, here we have limited ourselves to three of the most common ones, namely: Gaussian, log-normal and Poisson. It is easy to see that the state estimation problem for the Gaussian distribution, if variances are treated as uniform, is identical to the Non-Negative Matrix Factorization (NMF) problem. Thus, we utilize standard NMF solvers for this distribution followed by the column normalization of *W* as stated above. Similarly, for log-normal data, transforming the data as *Y* = log(1+*X*) makes the transformed dataset Gaussian distributed and allow us to again use NMF solvers. It has been our observation that column normalizing the Gaussian and log-normal distributed datasets before state estimation can lead to an improvement in results and hence has been performed in this paper.

### 2.2 Distribution selection

In our program we have a set of possible distributions to choose from. In order to select the distribution to use automatically, we implemented a method using fit error (Fig 2A). First, we fit the different distributions for each gene using maximum likelihood. Then, we compute the distance between the empirical distribution and each of the fitted distributions. This test is similar to the root-mean-square statistic for goodness of fit (Perkins *et al.*, 2011). The distribution with the minimum such distance is considered the best fit. Finally, we output the best estimation fraction vector which captures the fraction of genes for which each distribution is the best-fit distribution. This is meant to be a guideline for the selection of the sampling distribution during the matrix factorization process.

**Fig. 2.**
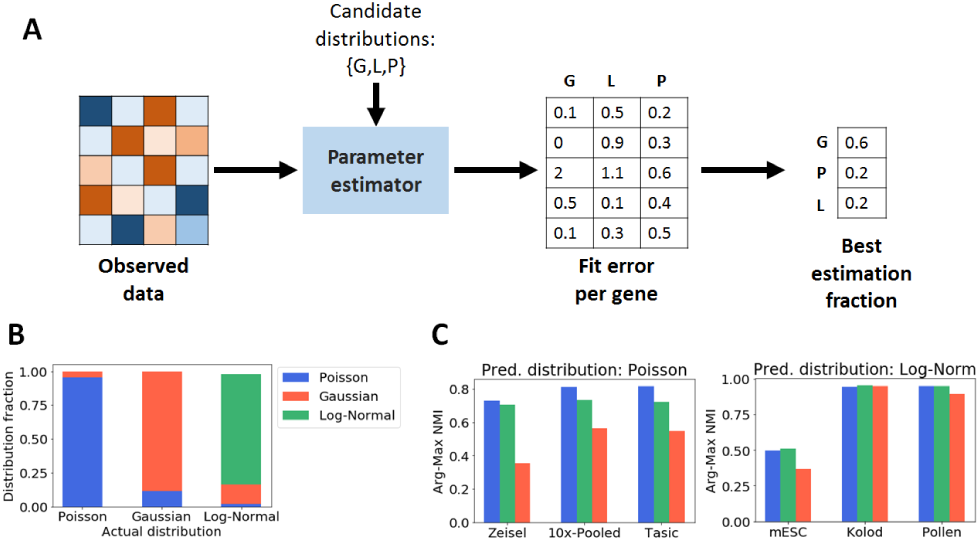
Selecting the best sampling distribution for a dataset from a set of distributions using Distribution Selector. A) Overview of Distribution Selector. B) ‘Best Estimation Fraction’ correctly identifies distributions of synthetic datasets. C) Comparison on different single cell datasets show that using predicted distribution leads to the highest cluster purity (measured using arg-max NMI).

### 2.3 Initialization for state estimation

Since state estimation is a non-convex problem, its result depends greatly on the initialization. Two commonly used methods for NMF initialization are based on k-means and SVD (singular value decomposition), respectively (Langville *et al.*, 2006; Boutsidis and Gallopoulos, 2008). In UNCURL, we have two ways to initialize the state estimation: (1) distribution-specific k-means initialization (2) truncated SVD + k-means based initialization. We describe our distribution specific K-means in the rest of the section, particularly for the Poisson case.

In the following sub-sections we develop an explicit framework to utilize prior biological information to initialize the matrix factorization. This relies on clustering each gene into clusters of high and low expression, which is done using k-means clustering for Gaussian and log-normal (after log-transformation) distributions. Here we outline a similar procedure for the Poisson distribution that allows our approach to be consistent across different distributions along with an approach to initialize the clustering.

#### 2.3.1 Poisson k-means++

k-means++ (Arthur and Vassilvitskii, 2007) is a well known seeding method for the k-means clustering algorithm, which tries to identify *k* points in the data with the highest mutual separation. While popular, its use of Euclidean distances between points makes it a poor fit for non-Gaussian distributed data. To use a similar approach for Poisson sampled data, we use a new distance metric. One intuitive distance would be the Poisson log-likelihood with one data point as the mean (say *y*) and the other as the point being considered (say *x*). Let (*ll*_*p*_(*x | y*) be defined as the log likelihood of a Poisson distribution with mean *y*. However, *ll*_*p*_(*x | y*) is not symmetric, which is required for a distance measure. We create a normalized version of this function which satisfies all properties of a distance measure:

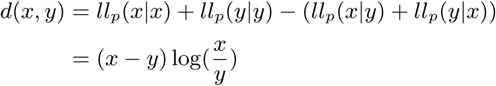

This distance is based on the observation that value of *ll*_*p*_(*x | y*) is maximum when *x* = *y*. Thus, the *d*(*x, y*) quantity measures the distance from the maximum value log-likelihood value for both *x* and *y* (for the sake of symmetry). This distance then replaces the Euclidean distance used in the standard implementation of k-means++.

#### 2.3.2 Poisson k-means

Poisson k-means is very similar to the classical k-means clustering with the difference being in the underlying distribution of the data; it is essentially the hard EM algorithm applied with a Poisson distribution. As with state estimation, we assume we are provided the expected number of cell types *k* and the data matrix *X ∈* ℝ^*n×d*^. The first step of the algorithm involves calculating the Poisson log-likelihood for each cell given a set of means (*M ∈* ℝ^*n×k*^), representing *k* cell types and then assigning each cell to the cell type for which it has the highest log-likelihood. The logarithm of the probability that cell *c* with observed counts *X*_*.c*_ is sampled from the type *i* with gene expression *M*_*.i*_ is then *ll*_*p*_(*c | i*) = − ∑_*j*_[*M*]_*ji*_ + [*X*]_*jc*_ log([*M*]_*ji*_).

This is called the E step of the algorithm, which partitions the cells into distinct types. This is followed by the M step, where we find the means for each cell type that maximize the log-likelihood. For the case of the Poisson distribution, this is simply the arithmetic mean of the data given by:

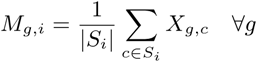

Here, *S*_*i*_ is the set of cell indices for which the log-likelihood is highest for the *i*th mean. The M step creates a new estimate of the means, which are then used to redo the E step. This procedure is repeated till convergence or until a maximum number of iterations.

### 2.4 Qualitative semi–supervision with QualNorm

In many scenarios where scRNA-seq is carried out, there is a wealth of prior knowledge. For example, there may be FISH images or bulk gene expression data measured through microarray or RNA-seq. Alternatively, marker genes may be known for a subset of cell-types. Two major issues in using such information are the incompatibility between different data types (e.g. FISH images or microarray data with RNAseq data), and variability between experiments using the same technique (e.g. bulk RNA-seq batch effects). We develop a method to specifically account for such variations in order to leverage this prior information. A key benefit with our method is that since the prior information is leveraged in UNCURL preprocessing, it can boost the performance of downstream methods not designed to utilize such prior information.

A basic problem that we need to solve is in deciding how to incorporate such differing type of information into our framework. Our hypothesis is that even though the particular quantitative measurements may not transfer well to scRNA-seq, the qualitative information inherent in such a dataset can be exploited. To do that, we first convert the available prior information into a binary matrix, that can then be imported into our method. In some cases, the prior biological knowledge available may be marker information which is directly in the binary format. In other cases, where prior biological knowledge is available as bulk gene expression or other real-valued measurements, we first binarize the gene expression by thresholding around the central value for each differentially expressed gene. In our experiments, we have used the Poisson version of t-test (Gu *et al.*, 2008) to identify genes that are composed of two separate distributions and have limited the input to only include up to 25 ‘ON’ genes per cell type with the highest p-values. Details of this process are described in the Supplementary Methods.

The inputs to the framework are the following: 1) A single cell sequenced data matrix *X ∈* ℝ^*n×d*^, 2) the number of cell types expected in the data, *k* and 3) a binary matrix of dimension 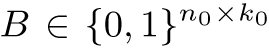, where *n*_0_ is the number of genes for which the information is provided and *k*_0_ is the number of cell types for which the information is provided. We note that the number of cell types *k*_0_ for which prior information is available can be lesser than the number of cell-types *k*, and the number of genes *n*_0_ for which prior information is available can be lesser than total number of genes in the data *n*.

Now, the main algorithmic problem is how to utilize the matrix *B* in solving the state-estimation problem. The obvious approach is to utilize the matrix as an initialization for state estimation. However there are three bottlenecks in our problem. (1) The matrix is binary, not real valued and it is unclear how to utilize such a matrix. (2) Information about every cell type is not available, i.e., *k*_*o*_ *≤ k*. (3) Information is not available about every gene, i.e., *n*_*o*_ *≤ n*. We deal with these issues sequentially.

Suppose we have a binary prior information matrix *B* such that *n*_0_ = *n* and *k*_0_ = *k*. We wish to convert this into a real valued matrix *M*_0_ of the same size, where the expression is in the same scale as the scRNA-seq data *X*. To do this, we use the data matrix *X* to calculate the high and low levels for each gene by clustering the observed expression values for that gene into two clusters and mapping the high and low values of cluster centers to 1 and 0 respectively. We note that in the case that *n*_0_ *≤ n* and *k*_0_ *≤ k*, still the same method can be utilized to obtain the real-valued matrix *M*_0_ of size *n*_0_ *× k*_0_.

Next, we take up the issue that information is not available about all genes, i.e., *n*_0_ *< n*. The basic idea that we exploit is the following: the knowledge even of some genes is sufficient to cluster all observed cells into *k*_0_ types, using a distribution specific clustering (for example, the Poisson k-means algorithm described earlier). The *k*_0_ cluster centers in dimension *n* can then be used as the means of these clusters, thus effectively giving us an updated *M*_0_ matrix of size *n × k*_0_.

Finally, we proceed to the issue that only some cell-types are specified, i.e., *k*_0_ *≤ k*. In case that *k*_0_ = *k*, we have an initialization for all the matrix *M*_0_, which can be used an initialization for the alternating maximization algorithm. In case that *k*_0_ = *k*, we do one round of distribution specific k-means++ with the first *k*_0_ components initialized by the known data and the rest obtained as maximally distant points from the known points. This returns us the means for all the *k* components, which we update as *M*_0_ of dimension *n × k*. This matrix of predicted means *M*_0_ is now used to initialize the various downstream algorithms, in particular serving as an initialization for maximizing *W* in the alternating optimization.

## 3 RESULTS

### 3.1 Distribution selector correctly predicts the best sampling distribution for a dataset

To verify the accuracy of the distribution selection methodology, we first generated three synthetic datasets using different distributions (Poisson, Gaussian and log-normal). Each gene in the synthetic dataset has a mean that was randomly chosen between 0 and 1, with a constant variance for all genes for the Gaussian and log-normally distributed datasets.

As seen in Fig 2B, distribution selector correctly predicts the dominant distribution for each of the synthetic datasets. On the real datasets, we do not know the underlying distribution, however, we can check which distribution performs best on the downstream task of clustering. We check whether the predicted distribution leads to the highest accuracy. The predicted clusters are identified by assigning each cell to the highest weight class in the cell type fraction matrix, W. The cluster purity is then measured using Normalized Mutual Information (NMI) between the predicted clustering and the true cell types in the data and is seen to be highest for the distributions that are predicted to be dominant distribution according to distribution selector as seen in Fig 2C. In general, the Poisson distribution is seen to be a better fit for count or UMI data, while the log-normal distribution is a better fit for normalized (FPKM, RPKM, TPM) data.

Having identified the sampling distribution for a dataset, the state estimation procedure factorizes the data into two matrices, *M* and *W* using the distribution, as described in 2.1.1. The product of these factorized matrices then can be treated as an estimate of the true state matrix and used for all subsequent downstream learning tasks.

### 3.2 Preprocessing leads to improvements in clustering and visualization

Clustering and dimensionality reduction are common downstream tasks for scRNA-seq data. These are both unsupervised tasks: clustering involves dividing the cells into different cell types based on similarity of gene expression patterns, while dimensionality reduction involves creating a low-dimensional view of the data for visualization or clustering. Both tasks are useful in identifying cell sub-populations in an unlabelled dataset.

Two of the most commonly used tools for scRNA-seq data are PCA and tSNE (Maaten and Hinton, 2008). While these algorithms have different underlying mechanisms for converting the high dimensional information to typically 2 or 3 dimensions, they assume Gaussian or t-distributions for the dataset, which is often not reflective of the actual sampling distribution of the data. To alleviate such issues, specialized tools have been developed for scRNA-seq data such as SIMLR (Wang *et al.*, 2017) and ZIFA (Pierson and Yau, 2015), which explicitly account for the distribution of scRNA-seq datasets. While ZIFA explicitly accounts for zero inflation, it does not account for the full sampling distribution. SIMLR uses multiple Gaussian kernels to fit the data without having an explicit sampling model. While these tools lead to improvements over PCA/tSNE, we demonstrate that using UNCURL as a preprocessing tool can greatly improve the performances of these common dimensionality reduction tools and often even make them better than specialized tools such as ZIFA/SIMLR.

UNCURL can be used for clustering in several ways. The simplest clustering method is to assign as the cluster *c*_*i*_ = argmax_*j*_ *W*_*j,i*_ for cell *i*, where *W* is inferred from state estimation. We call this method UNCURL-W. In addition, clustering can benefit from UNCURL preprocessing by running tSNE or PCA and then k-means on the output of UNCURL.

We demonstrate the utility of using UNCURL as a preprocessing tool by comparing the clustering performance of various methods with and without preprocessing with UNCURL, along with clustering after dimensionality reduction with ZIFA and SIMLR. Additionally we compare with another preprocessing tool, Magic (Dijk *et al.*, 2017). Performance was measured using the NMI between true and predicted clusters with the selected preprocessing and clustering methods, as done in Wang *et al.* (2017). As seen in Fig 3A, UNCURL improves the performance of k-means and PCA on all four datasets (a more comprehensive set of results can be seen in Supplementary methods), and it improves the performance of tSNE on most datasets. Moreover, in all four datasets the top performing approach (showed with the red dotted line) is an UNCURL preprocessed approach.

**Fig. 3.**
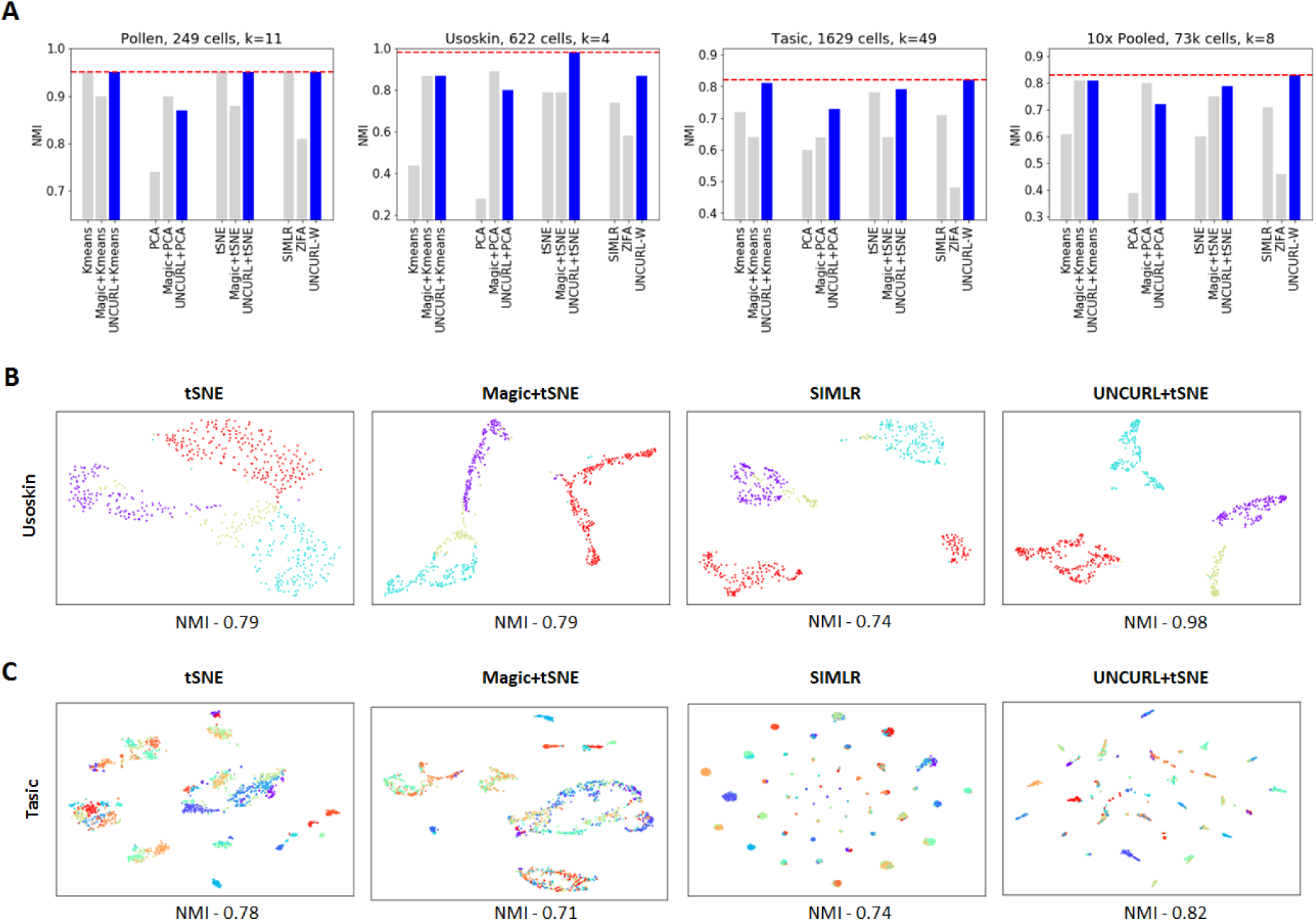
Preprocessing with UNCURL leads to improved visualization and clustering performances. A) Comparison of various clustering approaches with and without preprocessing on different scRNA-seq datasets. B-C) Different 2D visualizations of the Usoskin and Tasic datasets respectively.

To investigate the effect of UNCURL preprocessing on visualization, we run tSNE on the Usoskin *et al.* (2015) and the Tasic *et al.* (2016) datasets, which are FPKM and Count valued respectively. We compare against the unprocessed tSNE visualization, tSNE after preprocessing using Magic and visualization using SIMLR (which is closely related to tSNE). The Usoskin dataset (Fig 3B) comprises of sensory neuronal cells from four principal neuronal types. We see that while all other approaches end up grouping one or more cell types together, UNCURL leads to complete separation of all principal neuronal types.

The Tasic (Fig 3C) dataset is comprised of cells from mouse cortex tissue, with 49 cell types, including neuronal and non-neuronal cells. Using UNCURL along with tSNE, we were able to identify 49 distinct clusters that corresponded very strongly to the cell types identified previously. For the same dataset, the performance of tSNE without preprocessing and Magic preprocessing was seen to be significantly worse, with many cell types being grouped together.

### 3.3 Prior knowledge improves UNCURL

To demonstrate the utility of prior biological knowledge, we consider a dataset with available qualitative information in the form of bulk RNA-seq data obtained from different experimental conditions and consider the effect of semi-supervision on visualization with tSNE. Specifically, we focus on the a subset of the data set of Zeisel *et al.* (2015), comprised of five non-pyramidal cell types: oligodendrocytes, astrocytes, interneurons, microglia and endothelial cells. An upper bound on the performance with semi-supervision information is obtained when we feed the aggregate means of the true clusters (the means of all cells with each of the ground-truth labels) as the initialization. We compare this with semi-supervision using the output of QualNorm, bulk means and unsupervised preprocessing. In order to test the validity of our QualNorm framework, we compare the performance with aggregate-mean initialization to the performance obtained when we process these aggregate means through the QualNorm framework. In Fig 4B, the four initialization methods are compared, and it is seen that while semi-supervision with aggregate means and QualNorm means lead to a clear separation of cell types and an improved over unsupervised preprocessing, initialization with bulk means leads to worse visualization for this dataset. A similar experiment (see Supplementary methods) with the 10x pooled dataset also leads to a qualitative improvement in tSNE/PCA based visualizations and improvement in clustering purity.

**Fig. 4.**
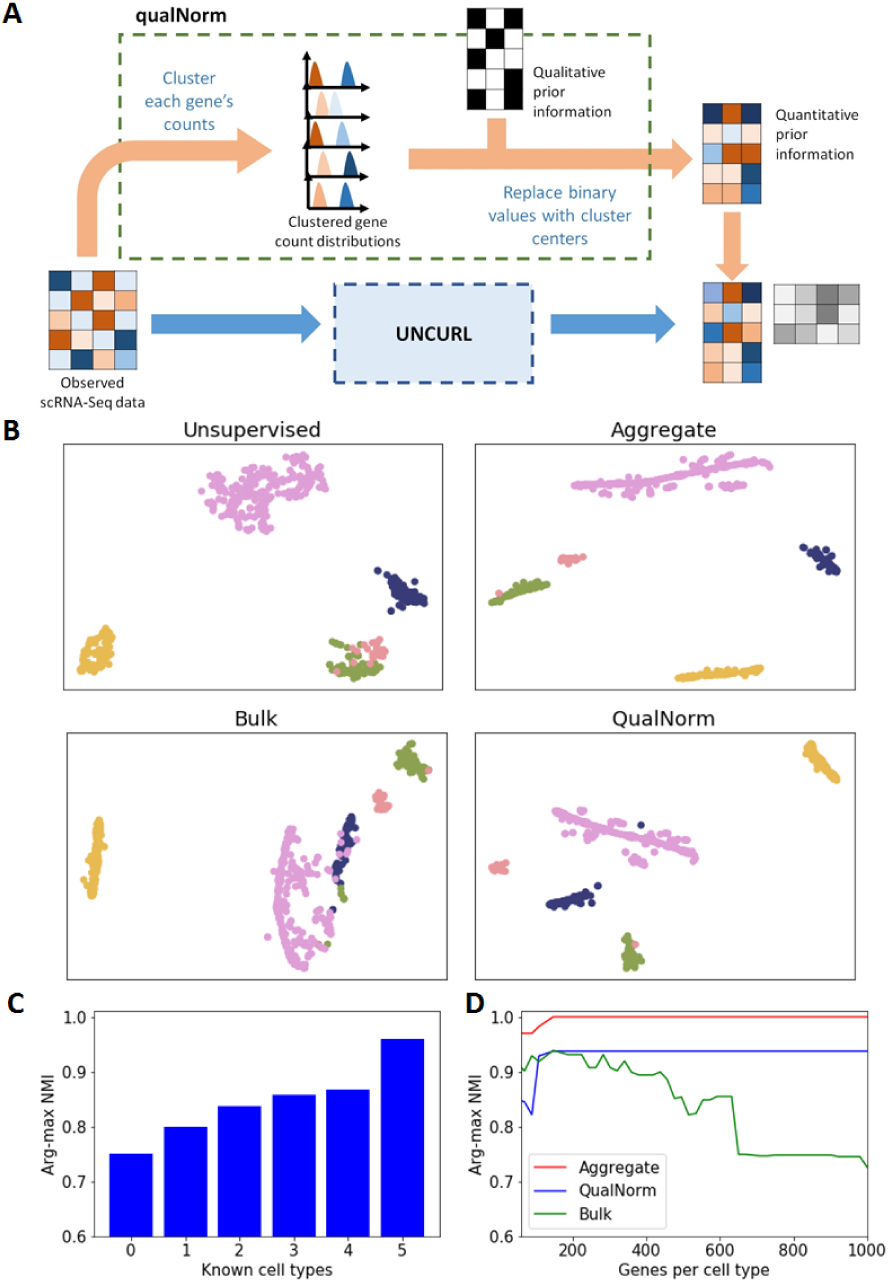
Semi-supervision with prior biological information can further improve the performance of UNCURL. A) An illustration of the qualNorm framework to convert qualitative prior information into good initialization points for unsupervised learning algorithms. B) Visualization using tSNE with unsupervised, aggregate, bulk and QualNorm semi-supervision on a subset of the Zeisel dataset (containing 1672 cells and 5 cell types). C) Comparison of improvement in purity with prior information about different number of cell types. D) Comparison of clustering purity with prior information about different number of cell type specific genes.

First, we simulate the scenario where we have information available about a subset of cell types by using different number of known means (generated by binarizing the public bulk data and passing it through the QualNorm) and generating the other means using our version of the distribution informed k-means++ algorithm. We then calculate NMI between predicted and true clusters (using arg-max of W) to quantitatively measure the performance of state estimation. As seen in Fig 4C, we observe that increasing the number of known means leads to improvement in accuracy. Moreover, we also see that prior information about even a subset of cell types is usually enough to improve the performance over the completely unsupervised case.

Then, to test the effect of having information about only a subset of the genes, we chose different number of top cell type specific genes (measured by one-vs-all differential expression) as initialization points for UNCURL and tested their effect on the purity of the clusters. Furthermore, we also tested the effectiveness of three different semi-supervision strategies namely, 1) true cluster centers (generated by taking aggregate means with the known true labels), 2) bulk means and 3) QualNorm means. It is seen in Fig 4D, that while the true cluster centers lead to almost perfect estimation of cluster membership, the QualNorm means lead to better accuracy than using the bulk data for most subset sizes, which we attribute to biases inherent to different sequencing methods. This effect becomes more pronounced as the number of genes being considered increases. An interesting observation here is that information about a few cell type specific genes is enough to very high NMI values, even when the information is qualitative.

While the results in these two cases of missing information highlight the flexibility of the QualNorm framework to handle different amounts of provided prior information, they also demonstrate how having even a little additional information is enough to improve unsupervised learning results significantly.

### 3.4 Improving lineage estimation with UNCURL

One biologically important downstream task post dimensionality reduction is lineage estimation, which is often used to study the dynamics of various genes during cell differentiation or development. Lineage inference aims at identifying smooth continuous manifolds in which cells lie in order to study the gradual change in gene expression during various developmental processes. While there exist several common tools that exist for this purpose (Trapnell *et al.*, 2014; Welch *et al.*, 2016; Qiu *et al.*, 2017), most tools operate directly on the sampled observed data. Additionally, most lineage estimation tools do not allow for the incorporation of any prior biological data beyond selecting the subset of genes to use for lineage inference. Here we demonstrate that the use of UNCURL preprocessing can lead to the estimate of cleaner lineages as well as allow for the incorporation of qualitative prior information into the lineage inference process.

Since it is not possible to obtain ground truth ordering of cells, we first study the effect of UNCURL preprocessing on simulated datasets. We created two separate synthetic lineages (a linear and a branched) using a method described in Supplementary Methods and sampled using a Poisson distribution. For both datasets, we preprocessed the datasets using UNCURL and Magic and tested three commonly used lineage inference tools (Monocle, Monocle2 and Slicer) on both preprocessed and unprocessed datasets (see Supplementary Methods). For the linear dataset, a good measure of lineage accuracy is the rank correlation of the true ordering of the cells with the pseudotime (an arbitrary metric that is commonly used to measure progress along a trajectory). We find that UNCURL improved the rank correlation of all three methods compared to both the unprocessed data and Magic, whereas preprocessing using Magic lead to worse performance for both Monocle2 and Slicer. For the branched data, there isnt a similarly simple way to quantify lineage estimates so we looked at the branch purity (a measure of how well cells have been ordered into the right branches). Even in this case we found that the branch purity obtained in Monocle2 after preprocessing with UNCURL was higher than both unprocessed and preprocessing using Magic.

We now look at real biological data with some amount of ground truth data available. We specifically focus on the dataset of Hanchate *et al.* (2015) which contains cell types comprised of four stages of olfactory neurogenesis along a linear differentiation path. The cell types were identified using markers specific to different stages of development. While this information itself is of the form of qualitative prior information, using the marker genes used to label the cells would make the problem trivial for UNCURL. Hence, we instead simulate a bulk dataset by using the true labels of the dataset, which is then binarized to generate qualitative marker information for all sufficiently expressed genes (same as used in the original paper). We then use this to use both the qualNorm initialization as well as unsupervised initialization to preprocess the dataset using UNCURL. We then perform lineage inference using Monocle2, Monocle and Slicer on the unprocessed data, UNCURL preprocessed data and Magic preprocessed data. As seen in Fig 5 A-C, QualNorm initialized UNCURL leads to consistent improvements of the lineage inference algorithms when measured by the amount of overlap between the known cell types in the lineage graphs. In addition, preprocessing using UNCURL without semi-supervision leads to qualitatively similar lineages as the unprocessed datasets with slight overlap between consecutive cell types (see Supplementary Methods). On the other hand, while preprocessing using Magic seems to lead to qualitatively similar lineages for Monocle2 and Slicer, it also leads to the estimation of a major non-existent branch in the case of Monocle (Fig 5B).

**Fig. 5.**
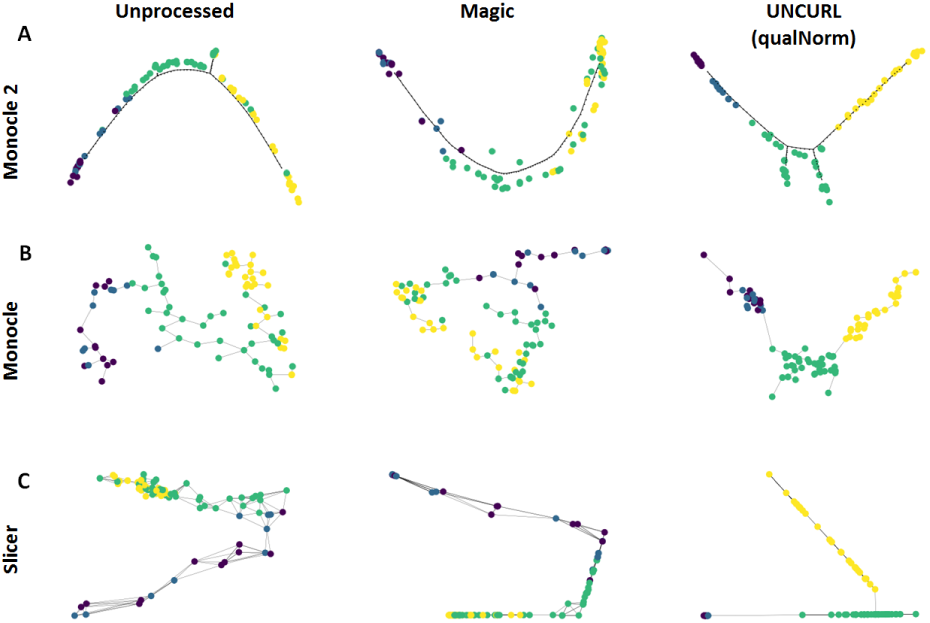
Semi-supervised preprocessing using UNCURL can dramatically improve the performance of lineage inference algorithms. A-C) Lineages inferred by Monocle2, Monocle and Slicer respectively with no-preprocessing, Magic preprocessed and semi-supervised UNCURL preprocessed for the dataset of Hanchate et. al. (containing 85 cells from 4 cell types). The lineages obtained after preprocessing using semi-supervised UNCURL lead to much clearer separation of known cell types in the predicted lineages.

### 3.5 Scalability

UNCURL is capable of running on larger datasets comprising of up to millions of cells. UNCURL uses a fast public NMF package (Pedregosa *et al.*, 2011) for the log-normal and Gaussian distributions, while using SPNoLips (Sparse-Parallel-NoLips, described in supplementary methods), which is a custom implementation based on the NoLips algorithm (Bauschke *et al.* (2016)) for the Poisson distribution. Both implementations are capable of using sparse matrices as input for memory and runtime advantages, and are parallelizable.

The runtime of UNCURL is *O*(*n*_*p*_*k*), where *n*_*p*_ is the number of nonzero elements in the input matrix *X*, and *k* is the number of cell types. This means that UNCURL scales linearly in the number of cells in the dataset. To deal with the dependence on *k*, one possibility is to use UNCURL hierarchically: first run it with a small *k* on the entire dataset, then partition the dataset based on the assigned clusters, and run UNCURL on the subsets.

The computational performance of UNCURL on various datasets compares favorably to that of other methods as seen in Figure 6. The runtimes of UNCURL is usually less than the other comparable methods for clustering tasks on most datasets as seen in Figure 6 (a more comprehensive comparison is in the supplementary methods). The memory usage was lower than that of comparable methods such as SIMLR and Magic. UNCURL’s performance is best on sparser datasets, where more entries are zero, since the NoLips update function only uses nonzero values of the data matrix. Runtime comparisons with Magic and ZIFA are limited because Magic’s memory usage is quadratic in the number of cells and ZIFA is slow compared to other algorithms, making it impractical to run on the largest datasets.

**Fig. 6.**
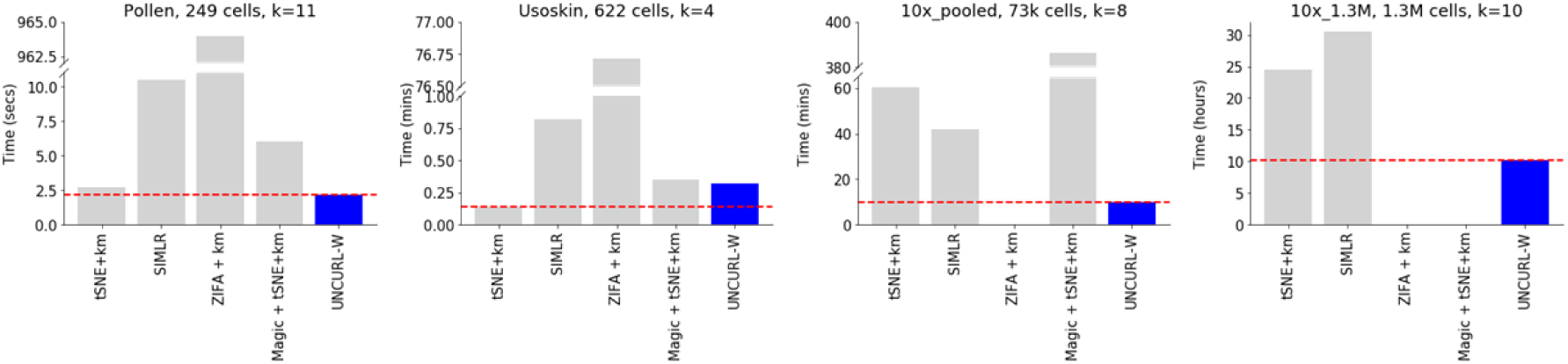
Timing comparison of different clustering approaches for various scRNA-seq datasets. UNCURL is faster than other approaches on most datasets of various sizes. Moreover, unlike methods like ZIFA and Magic, UNCURL is scalable for large datasets. A more comprehensive comparison can be found in the supplementary methods.

### 3.6 Exploratory analysis of the 10x 1.3 million dataset

The scalability of UNCURL allows it to run on very large data matrices, including the 1.3 million-cell dataset from 10XGenomics (2017). This dataset is composed of unsorted brain cells from 18-day mouse embryos. We tested UNCURL on both the full dataset and a 20,000 cell subset. Since this dataset does not have any ground truth labels, we used various exploratory methods to characterize the different cell types present.

We empirically chose the number of main cell types to be 10 after experimenting with various values of k. Our selection of k was based on the distinctness of genetic signature of the identified clusters. Fig 7A show the result of UNCURL’s clustering and visualization on the 20,000 cell subset while 7B shows the unprocessed tSNE visualization followed by k-means clustering. To test the concurrency of the two approaches, we generated a confusion matrix (Fig 7D) and noticed decent overlap between the clusters resulting from the two distinct methods (with an NMI of 0.45).

**Fig. 7.**
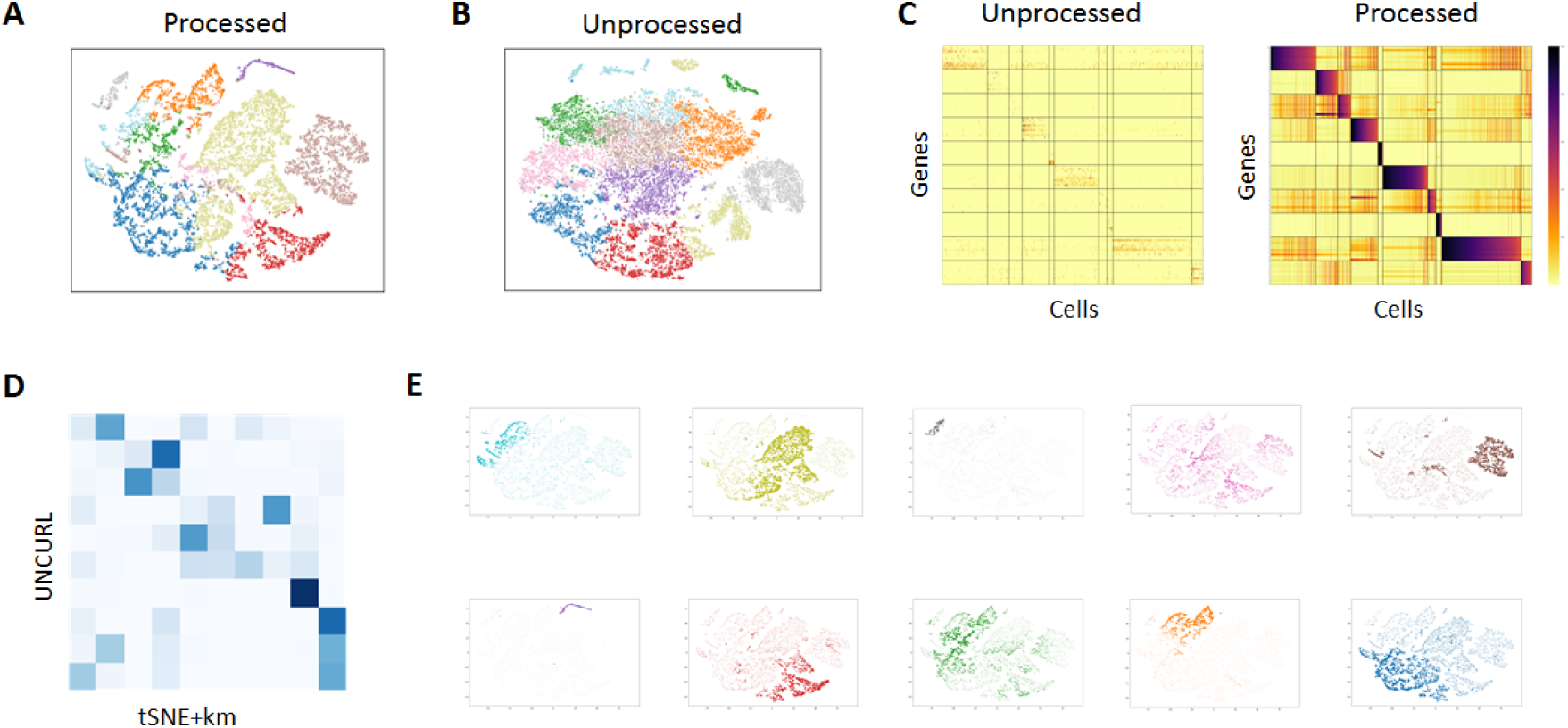
Exploratory analysis of the 10x 1.3 million cell dataset with UNCURL. A) tSNE plot on UNCURL preprocessed data with argmax inferred labels. B) tSNE plot without preprocessing with k-means inferred labels. C) Clustered heatmaps showing the top cluster specific genes identified by UNCURL before and after preprocessing. Cells sorted by decreasing W for each cluster. The heatmaps demonstrate that UNCURL identifies distinct sub-populations of cells and preprocessing makes the expression of the top clusters more distinct. D) Confusion matrix between UNCURL and tSNE + k-means labels. E) Average expression of the top cluster specific genes overlaid on the UNCURL processed tSNE plot. The expression for each cluster is colored to correspond the coloring used in A. It can be seen that the average expression of the top genes are very cluster specific, indicating that they identified distinct sub-populations.

To evaluate the clusters generated by UNCURL, we identified the genes most highly expressed in each cluster compared to other clusters (see supplementary method for description). We then created a subset of the expression matrix, where the rows are grouped by cluster-specfic genes and columns are grouped by cluster-specific cells. Visualizing the expression heatmaps before and after UNCURL (Fig 7C) shows that the top cluster-specific genes are distinctly expressed only in their individual clusters. Moreover, this expression pattern is amplified in the UNCURL processed data compared to the unprocessed data. This pattern is not seen when the data is preprocessed with Magic (see supplementary methods). To further validate our findings, we overlay the average expression of the top cluster-specific genes for each cluster on the tSNE visualization (Fig 7E).

## 4 CONCLUSION

In this manuscript, we introduced a preprocessing framework for scRNA-seq data. Our framework, UNCURL, uses the estimated sampling distribution of scRNA-seq data together with a convex mixture model assumption to estimate a true state matrix from observed scRNA-seq data. UNCURL further includes a computational framework, qualNorm, which can be used to incorporate prior biological knowledge into an improved estimate of the true state matrix.

By comparing against several benchmarking datasets, we demonstrated that preprocessing using UNCURL leads to superior separation of cell types in reduced dimensions as well as higher cluster purity for clustering tasks compared to prior tools. We further showed that semi-supervision using different types of prior information can lead to further improvement in accuracy of the learning tasks. Furthermore, we demonstrate that semi-supervised preprocessing using UNCURL allows the incorporation of prior information in even lineage estimation tasks. UNCURL scales to large datasets and typically runs faster than prior methods, particularly on large and sparse datasets. The run time for UNCURL scales linearly with the number of cell types in the dataset, but it may be possible to further reduce run-time using a hierarchical strategy.

UNCURL is an efficient preprocessing framework for several unsupervised and semi-supervised learning tasks, but it still has some limitations. While our method accounts for the sampling effect on the data, we do not take into account other sources of variability such as cell cycle effects and biological noise (Barron and Li, 2016). Moreover, presently the semi-supervision framework can only process prior information that can be converted to a binary format. While this still leads to improvement in accuracy, not all genes have binary states. Future work will be aimed at developing a learning framework that account for these other sources of variability and a more inclusive semi-supervision framework. We also plan to expand the number of sampling distributions available in the current software package to include more potential sampling distributions.

